# Label-free non-invasive quantitative measurement of lipid contents in individual microalgal cells using refractive index tomography

**DOI:** 10.1101/226480

**Authors:** JaeHwang Jung, Seong-Joo Hong, Han-Byeol Kim, Geon Kim, Moosung Lee, Seungwoo Shin, SangYun Lee, Dong-Jin Kim, Choul-Gyun Lee, YongKeun Park

## Abstract

Microalgae are promising candidates for biofuel production due to their high lipid content. To facilitate utilization of the microalgae for biofuel, rapid quantification of the lipid contents in microalgae is necessary. However, conventional methods based on the chemical extraction of lipids require a time-consuming destructive extraction process. Here, we demonstrate label-free, non-invasive, rapid quantification of the lipid contents in individual micro-algal cells measuring the three-dimensional refractive index tomograms. We measure three-dimensional refractive index distributions within *Nannochloropsis oculata* cells and find that lipid droplets are identifiable in tomograms by their high refractive index. In addition, we alter *N. oculata* under nitrogen deficiency by measuring the volume, lipid weight, and dry cell weight of individual cells. Characterization of individual cells allows correlative analysis between the lipid content and size of individual cells.

---

Biofuels have attracted increasing attention as sustainable energy resources that could potentially replace fossil fuels. Among the candidates suggested for biofuel production are a variety of plants^1, 2^, fungi^3^, bacteria^4, 5^, and microalgae^6^. The latter has long been considered a promising sustainable resource due to their rapid growth and high lipid content^7, 8^. Extensive research has been conducted to find suitable species and to optimize cultivation conditions to make large-scale lipid production by microalgae economically feasible^9, 10^. Even after this attention, there is still a high demand for development of new strains and optimized cultivation processes to facilitate industrialization of microalgae-based biofuels.

To enhance the research and development of microalgae strains for biofuel applications, it is essential to develop rapid, quantitative methods for lipid detection. Conventionally, gravimetric methods and chromatography have been widely used to quantify lipid content in microalgae. Because these methods are based on chemical extraction of the lipids using organic solvents; however, the methods require laborious, time-consuming extraction processes that take from several hours to days. Furthermore, the destructive nature of the extraction leads to irreversible loss of a large volume of each sample, which restricts repeated measurements. As an alternative, lipid quantification based on microscopic imaging techniques has recently been proposed in conjunction with advances in the relevant imaging techniques and instruments. Compared with quantification based on chemical extraction, quantification via imaging has the advantages of speed and low sample consumption. For example, fluorescence microscopy employing lipophilic fluorescent dyes is a representative imaging technique for visualizing and quantifying the lipid content in individual cells^11, 12^. Unfortunately, the fluorimetric methods are limited in principle to qualitative results because the permeability of cell membranes to dyes varies among microalgae species and the dye solutions used^12, 13^. In addition, invasive exogenous dyes may lead to incorrect results by affecting the physiology and viability of the cells. In contrast, Raman microscopy is a demonstrated quantitative, label-free imaging technique^14, 15, 16^. Raman microscopy detects molecular vibrational spectra, which are chemical fingerprints of molecules. Therefore, the lipid droplets should be specifically identifiable from their characteristic Raman spectral peaks. In spite of the outstanding molecular specificity of Raman microscopy, however, the intrinsically weak signals from Raman scattering require long acquisition time and a high-powered excitation light: both not desirable when investigating photosynthetic organisms.

Recently, optical diffraction tomography (ODT) or holotomography has emerged as a technique to provide quantitative three-dimensional (3D) imaging of biological samples without exogenous agents^17, 18, 19, 20^. As an optical analogy to X-ray computed tomography, ODT reconstructs the 3D refractive index (RI) distribution of a sample from multiple 2D holographic images of the sample obtained at various illumination angles. Because RI is an intrinsic optical property of materials, ODT provides morphological and biochemical information with sub-micrometer resolution^21^. Due to its capability for label-free, quantitative imaging, ODT has been verified as an imaging technique effective for studying biological samples including red blood cells^22, 23, 24^, white blood cells^25, 26^, yeast^27^, bacteria^28, 29, 30, 31^, phytoplankton^32^, and eukaryotic cells^33, 34, 35^.

In this paper, we present label-free, non-invasive, quantitative measurements of lipid content in individual micro-algal cells using ODT. By measuring 3D RI tomograms of living microalgae *(Nannochloropsis oculata)* we show that the lipid content in cells can be identified and quantified from the distinctively high RI values of the lipids. We also present results from our investigation of the morphological and compositional alteration of *N. oculata* under nitrogen deficiency (NDF), which triggers lipid accumulation in the cells. Volumes, dry cell weight (DCW), and lipid weight of individual cells are measured and compared with the results obtained with conventional methods. The capability for single-cell characterization allows correlative analysis between cell size and lipid content of individual micro-algal cells. The label-free quantitative imaging capability of ODT presents a promising potential breakthrough for industrial quantification of lipids, as well as providing a useful imaging technique for biological research to investigate the physiology of many kinds of living cells.

## Results

### Measurements of the 3D RI distributions within *N. oculata* cells

The 3D RI tomograms of *N. oculata* cells were produced employing ODT. The schematic diagram for the ODT measurements is presented in Fig. 2. While the cell was illuminated with a plane-wave laser beam at various incident angles (Fig. 2a), corresponding holograms were recorded using interferometry (Fig. 2b). The diffracted optical field information, which contains both amplitude and phase delay images, was quantitatively retrieved from individual holograms (Fig. 2c). Then the 3D RI distribution, *n*(*x*, *y*, *z*), of the cell was reconstructed from multiple 2D holograms (see Method for details). As shown in Fig. 2D and 2e, regions with distinctively high RI are found in the measured RI distribution of the cell. Considering that lipids are known to have a higher RI than protein^36, 37^, it was suspected that those high-RI regions were lipid droplets.

**Fig. 1.**
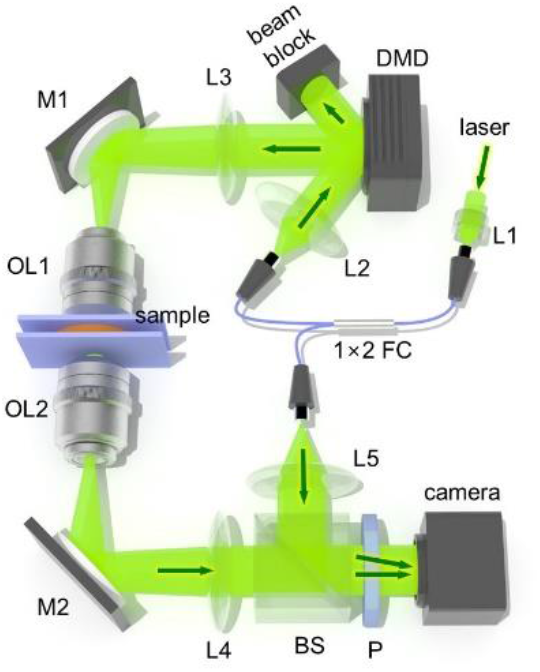
Optical system used for ODT: The optical system is based on Mach-Zehnder interferometric microscopy employing a digital micro mirror device (DMD). Light from a laser coupled in a fiber coupler is split into a sample beam and a reference beam. The sample beam illuminates a sample with incident angles controlled by patterns displayed on the DMD. A beam splitter in front of a camera constructs off-axis geometry for interferometry. This system has a maximum field of view of 50×50 μm^2^ with a total magnification of 60×. L1-5 lens; M1-2, mirror; FC, fiber coupler; OL1-2, objective lens; BS, beam splitter; P, polarizer.

**Fig. 2.**
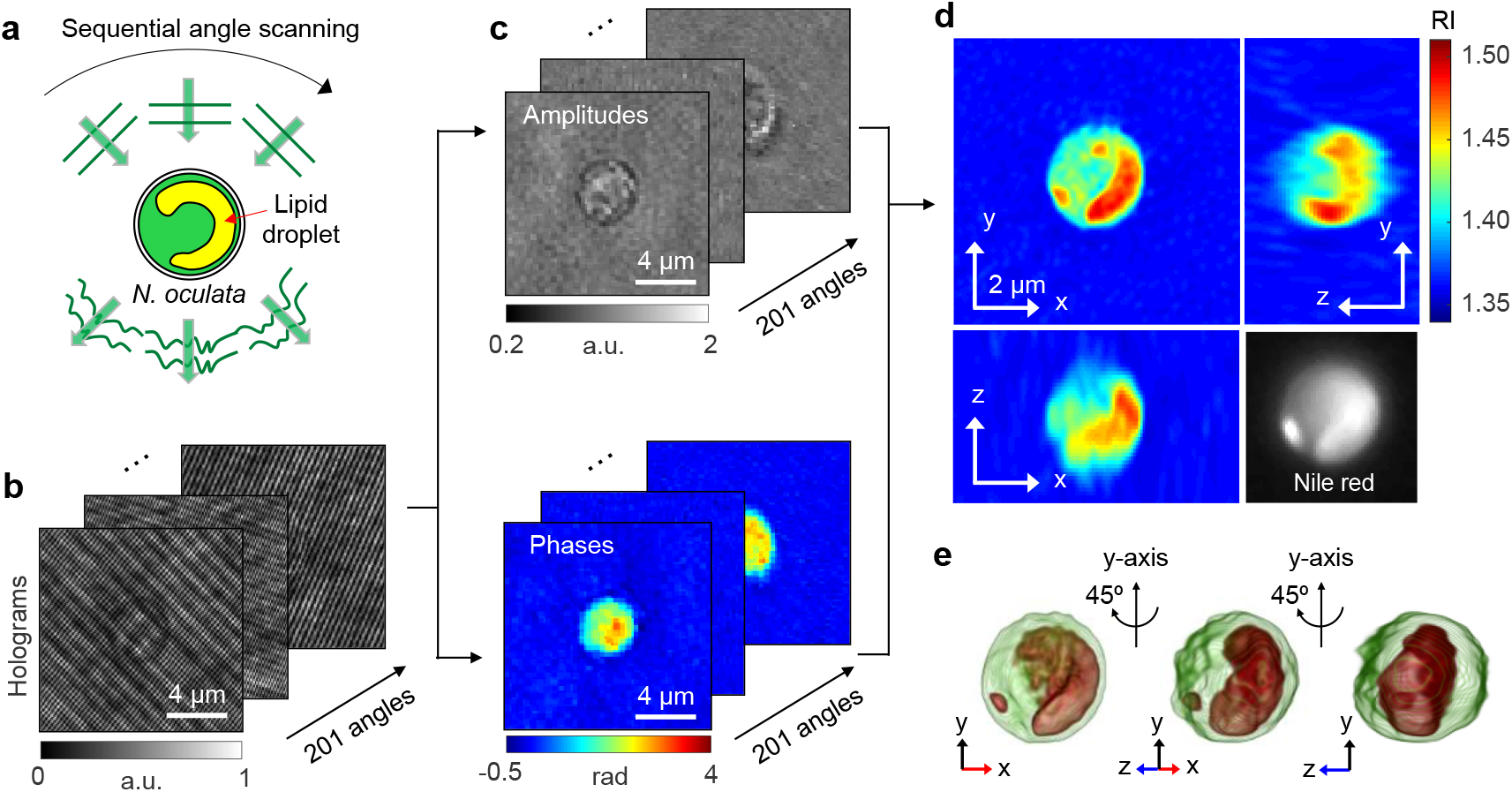
Schematic diagrams of the label-free identification of lipid droplets in individual *N. oculata* cells using ODT: (a) The sample is consecutively illuminated by a plane wave at various incident angles. (b) The holograms are recorded at 201 incident angles. (c) Retrieved amplitudes and phases of the optical fields diffracted by the sample. (d) Tomograms of the reconstructed 3D RI distribution of *N. oculata* in the x-y, y-z, and x-z planes. The Nile red fluorescence image of the same cell is shown in the lower right corner for comparison. (e) The 3D rendered iso-surface image of the reconstructed RI distribution at various viewing angles.

### Identification of lipids in the 3D RI distribution

To confirm whether the high RI regions in the 3D RI distribution were lipid droplets, the RI tomograms were compared with fluorescence images of the same cell labeled with Nile red. As shown in Fig. 3, there is coincidence between high RI regions in the RI tomograms (Fig. 3b) and fluorescence images (Fig. 3e) at multiple axial planes. For better comparison, we specifically identified the high RI regions (*n* > 1.46; Fig. 3c) that corresponded to the average RI of vegetable oils^38, 39^. Then we emulated fluorescence-like images from the masked RI tomograms by applying imaging processing based on 3D convolution (Fig. 3d). The comparison showed good agreement (average correlation 0.922) between the emulated images and the Nile red fluorescence at three axial planes (Fig. 3d and 3e). This result validates the notion that the threshold *n* > 1.46 is reasonable for the identification of lipid droplets.

**Fig. 3.**
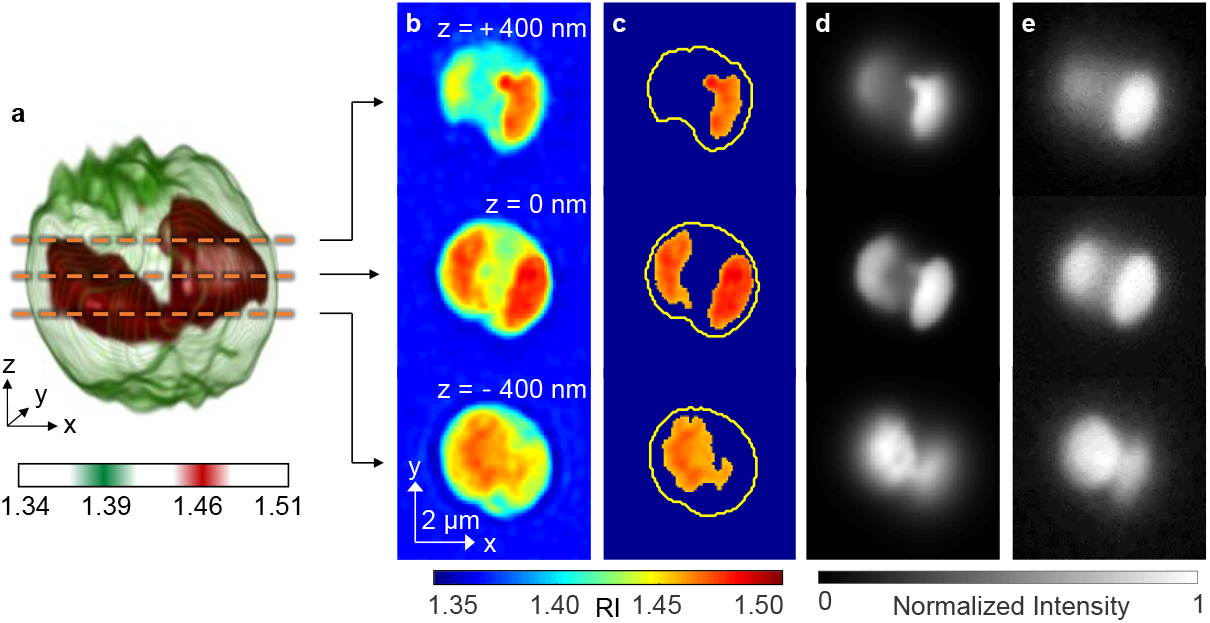
Identification of the lipid droplets inside an *N. oculata* cell: (a) A 3D rendered iso-surface of RI distribution of the cell. (b) Cross-sectional images of different axial planes. (c) The identified lipid regions based on the RI threshold *n* > 1.46. The yellow curves indicate the cell boundaries. (d) The emulated fluorescence-like images generated from the identified lipid regions. (e) The fluorescence images of the same cell stained with Nile red dye.

### Quantitative assessments of the effects of the nitrogen deficiency on individual *N. oculata*

To demonstrate the applicability of the present method, we investigated morphological and biochemical alterations in *N. oculata* cells caused by NDF. The 3D RI distributions of 42–53 cells were measured per group for four days. Because nitrogen has important metabolic functions in microalgae, the cells cultured under the NDF are expected to generate greater lipid content^40, 41^. The representative RI tomograms of *N. oculata* for the control and NDF groups for each day are shown in Fig. 4. Day 0 is defined as the moment after the cells were suspended in the nitrogen-depleted medium. From Day 0 to Day 3, the individual cells exhibited diverse intercellular RI distributions. In spite of the diverse morphology of the cells, however, the cells cultured under NDF generally have a larger volume of the lipids inside the cells, as shown in Fig. 4(h).

**Fig. 4.**
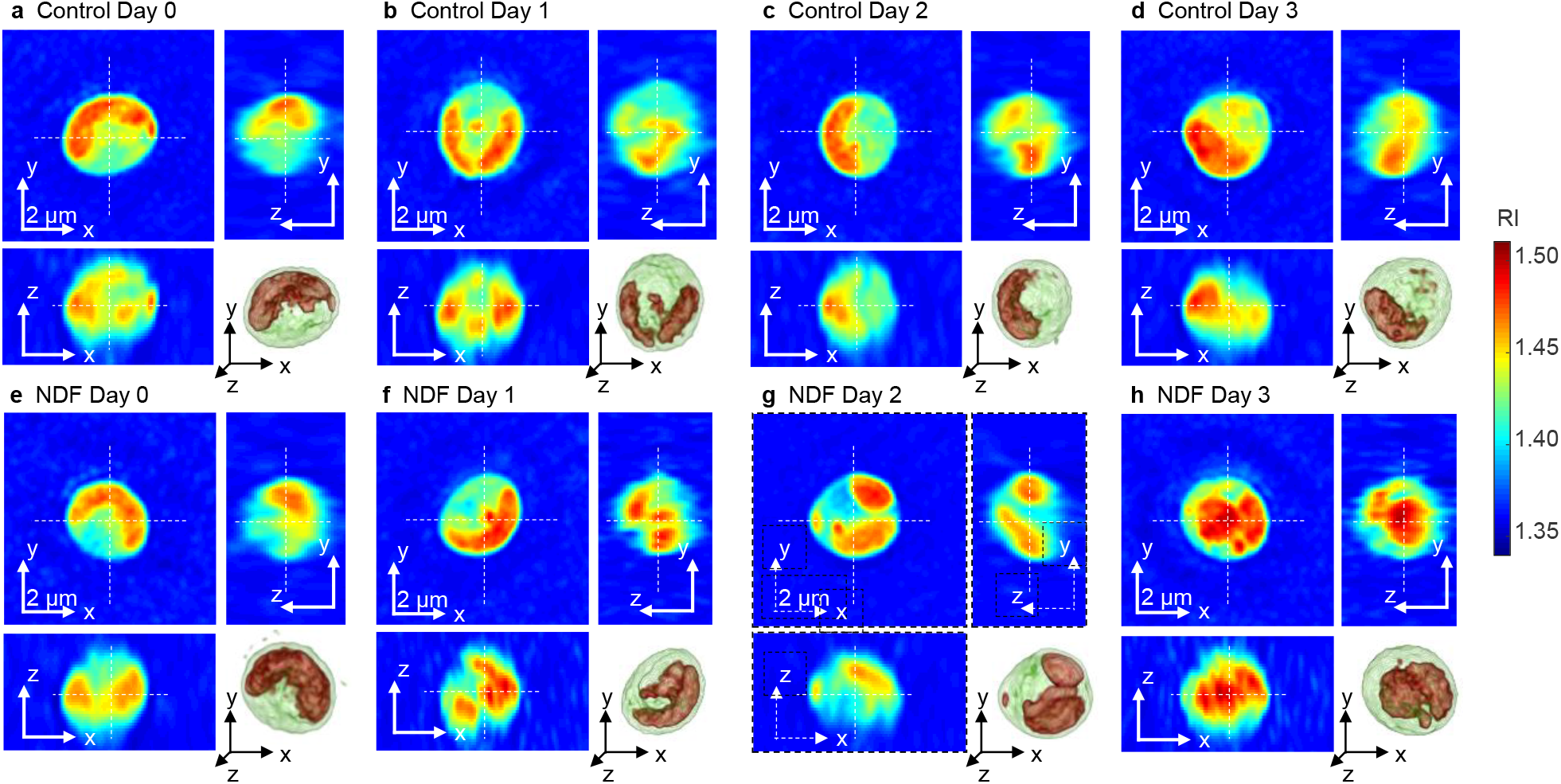
Representative tomograms of the 3D RI distributions of *N. oculata* cells in the control (a-d) and NDF (e-h) groups from Day 0-3, respectively. Corresponding 3D rendered images are shown in the lower right corner for each sample. The cells show diverse RI distributions.

In order to investigate quantitatively the lipid generation in cells and the effects of NDF, we calculated the cell volume, DCW, lipid weight, and lipid ratio of individual cells from the measured 3D RI distributions. The cellular volume was calculated from the cell boundary in the RI tomograms. The DCW was obtained as the summation of the weights of total cells, including the lipid and non-lipid components. The lipid weight inside the cell was obtained by multiplying the volume of the identified lipid regions (*n* > 1.46) by the average density of the vegetable oil (0.9 g/mL). The weight of the non-lipid components was calculated using the parameter called RI increment (RII), defined as an increment of an RI of a solution per an increment of a concentration of a solute^42, 43, 44^. Because the cell volume, except for the lipid regions, could be considered an aqueous solution with various components, the RI at non-lipid regions could be expressed as

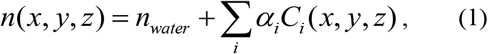

where *n*_water_ represents the RI of water, and α_*i*_ and *C_i_* indicate the RII and concentration of an arbitrary component i, respectively. Considering that one of the major non-lipid components inside the cell is protein, the average concentration of non-lipid biomolecules could be obtained by supposing the average RII is 0.185 mL/g, which is a typical value for proteins^42^. This RII value is also comparable to that of sugars^45^ and salts^46^, which are other major components in the cells. Therefore, by assuming the RII is constant for non-lipid components, the total concentration of the non-lipid components *C*(*x, y, z*) could be obtained using Eq. (1). Then integrating the concentration over the non-lipid volume would yield the weight of the non-lipid components. Finally, the lipid ratio was calculated as the ratio of the lipid weight to the DCW.

The cell volumes were measured to investigate the change in cell sizes. As shown in Fig. 5a, the results show that the cell volumes of both the control and NDF groups do not significantly change during cultivation. The constant cell volume of the control group is explained by the fact that the measurements were performed during the exponential phase when the cells actively divide. Meanwhile, the constant volume of the NDF group suggests suppressed cell growth due to the NDF. The measured volumes of *N. oculata* are mostly distributed within the range 10–20 fL, while the smallest and largest ones have volumes of approximately 5 and 33 fL, which volumes correspond to the equivalent diameter of 2.2 and 4.0 μm, respectively. Here, the equivalent diameter is defined as the diameter of a sphere of the same volume as the cell. The volumes obtained from the 3D RI distributions are consistent with independent measurements of volumes calculated from the diameter measured using a Coulter counter.

**Fig. 5.**
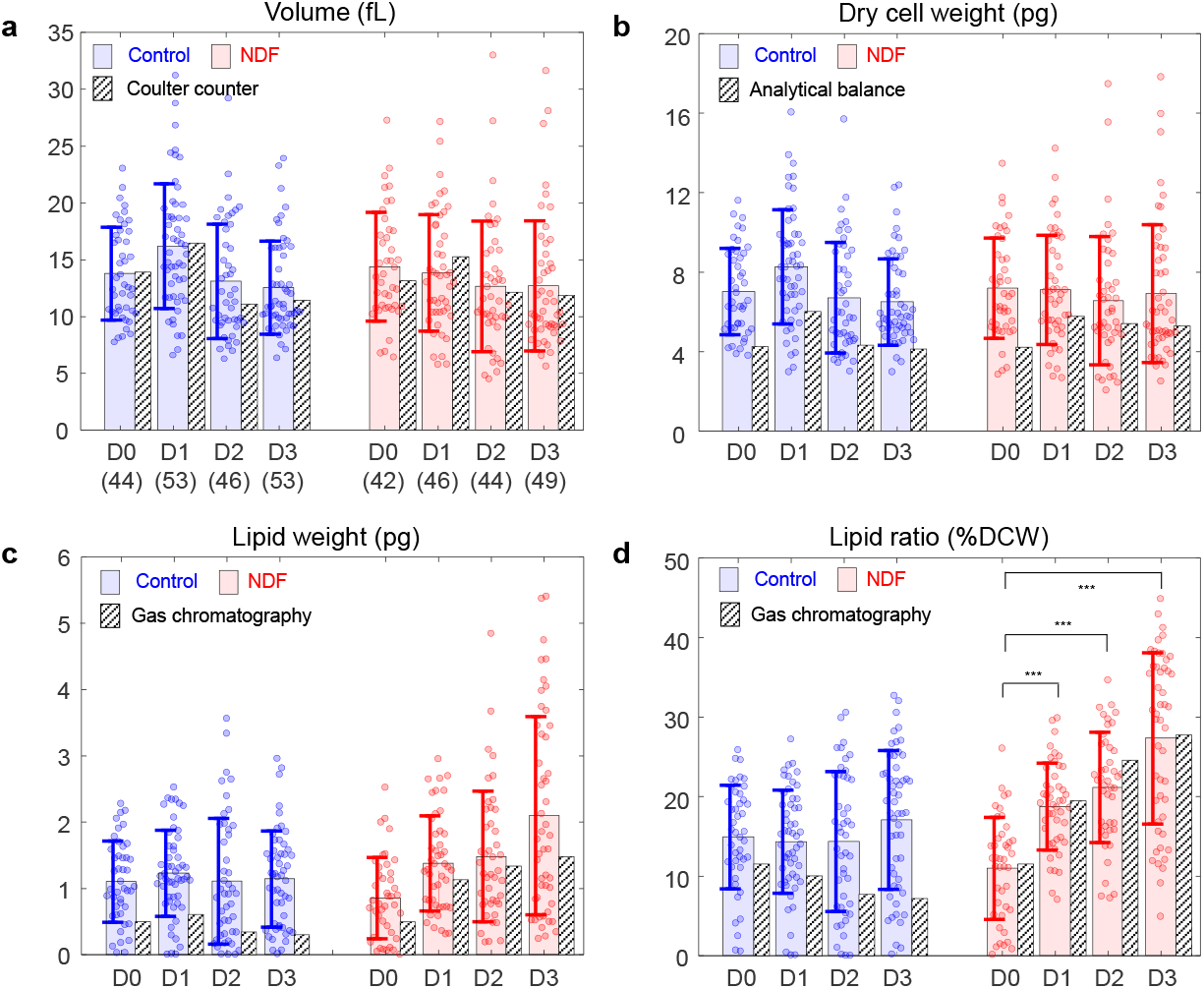
Measured (a) volume, (b) DCW, (c) lipid weight, and (d) lipid ratio of the individual *N. oculata* cells of the control and NDF groups. The numbers of the cells for each group are presented in parentheses. The circles represent the individual values measured by the ODT. The shaded color bars and error bars represent the mean values and standard deviations of the individual measurements. The hatched bars represent the values measured by conventional methods. Statistical tests presented in (d) were conducted using Student’s *t*-tests (***: *p* < 0.001). D0, D1, D2, and D3 indicate Day 0, 1, 2, and 3, respectively.

To investigate changes in the weight of total (all cell) components during cultivation under the NDF condition, the DCW of the cells was measured and analyzed (Fig. 5b). The measured DCWs of the individual cells in the control group are (7.03±2.17, 8.28±2.87, 6.72±2.79, and 6.51±2.17) pg (mean ± standard deviation) while the cells in the NDF group have DCWs of (7.19±2.52, 7.12±2.75, 6.57±3.22, and 6.93±3.47) pg from Day 0, 1, 2, and 3, respectively. To confirm the measurements of the dry cell weight using ODT, we also performed independent measurements using a gravimetric method and an analytical balance. Because the gravimetric method provides the bulk weight of the components in units of g/L rather than *per cell*, the bulk DCW was converted into the DCW per cell by dividing the DCW by the cell number density measured by the Coulter counter. Although the values measured using the ODT and the analytical balance show differences of up to 25%, they also show consistent tendencies. The discrepancy in these two measurement techniques may arise from at least two causes: i) the assumption that the non-lipid components have the same RII, and ii) the binary classification of cellular components as either lipids or non-lipids.

The measurements of the lipid weight were based on the assumption that the lipid accumulation inside the cell continuously increased under the NDF condition (Fig. 5c). The average lipid weight inside the cells increased 2.5 times during three days of the cultivation under NDF. The lipid weights of the cells in the NDF group are (0.85±0.61, 1.38±0.72, 1.48±0.98, and 2.10±1.49) pg from Day 0, 1, 2, and 3, respectively. In contrast, the algae in the control group show constant lipid weights of (1.10±0.61, 1.23±0.65, 1.11±0.95, and 1.14±0.72) pg from Day 0, 1, 2, and 3, respectively. This observation is consistent with the fact the lipid accumulation is induced by the nitrogen starvation^47, 48^.

Considering that the volume and DCW remained constant over four days, the increase in the lipid weight under the NDF implies that there should be some loss of other components such as proteins or carbohydrates during the lipid accumulation. This interpretation is consistent with previous reports that these microalgae decompose proteins to sustain metabolism when external nitrogen sources are depleted^49, 50^.

To validate the values retrieved from the ODT, the lipid weight in the bulk volume was also measured using gas chromatography and converted to the lipid weight per cell using the cell number density. The results from gas chromatography are qualitatively consistent with the ODT measurements. In the gas chromatography measurements, the average lipid weights of the cells in the NDF group increased (0.49, 1.12, 1.33, and 1.47 pg). The values obtained with gas chromatography were approximately 25% lower than the ODT results. One reason for this overestimation of the lipids in the ODT results might be due to the optical resolution. Because of the diffraction-limited optical resolution, the lipid droplets with size comparable or smaller than the optical resolution of ODT would appear slightly larger than actual volume. To reduce the effects of overestimation, more carefully designed RI threshold criteria for lipids should be introduced, for example, considering the spatial gradient of the RI distribution.

The lipid ratio in microalgae is generally utilized as a representative parameter for their lipid accumulation. From the measured RI tomograms, the lipid ratio was calculated from the measured DCW and lipid weight (Fig. 5d). The retrieved lipid ratios were (15.0±6.5, 14.3±6.5, 14.4±8.8, and 17.07±8.72)% for the control group and (11.0±6.43, 18.8±5.5, 21.2±6.9, and 27.34±10.77)% for the NDF group from Day 0, 1, 2, and 3, respectively. In contrast to the constant lipid ratio of the control group, the NDF group shows the lipid ratio increasing from 11 to 27% during the three days of culture. The increase of the lipid ratio is explained by the finding that the lipid weight increased while the DCW remained constant. The results obtained from the 3D RI tomograms show good agreement with the results measured by gas chromatography, especially for the NDF group.

### Correlative analysis of the properties of the individual cells

In addition to the investigation of the measured values for each culture condition, correlative analysis between the individual cellular properties would provide further insights into how the lipid weight and ratio change under NDF. Using conventional solvent-based methods that only measure the properties of bulk samples, access to the individual cellular properties was limited. In contrast, because the individual cellular characteristics could be measured using ODT, we could investigate the correlation between the lipid content and volume of individual cells (Fig. 6). The scatter plots of the cell volume versus lipid weights of individual cells present a positive correlation with coefficients of 0.65 and 0.76 (on average) for the control and NDF group, respectively (Fig. 6a and 6b). The positive correlation indicates that bigger cells have more lipids. In addition, the slopes of the linear fitting indicate that more lipids were accumulated (for similar cell volumes) after the cells were cultured under NDF. Considering the result that the cell sizes were not significantly changed during cultivation, these findings imply that the lipid productivity per cell is determined by the size of the cell at the moment that nitrogen depletion occurs.

**Fig. 6.**
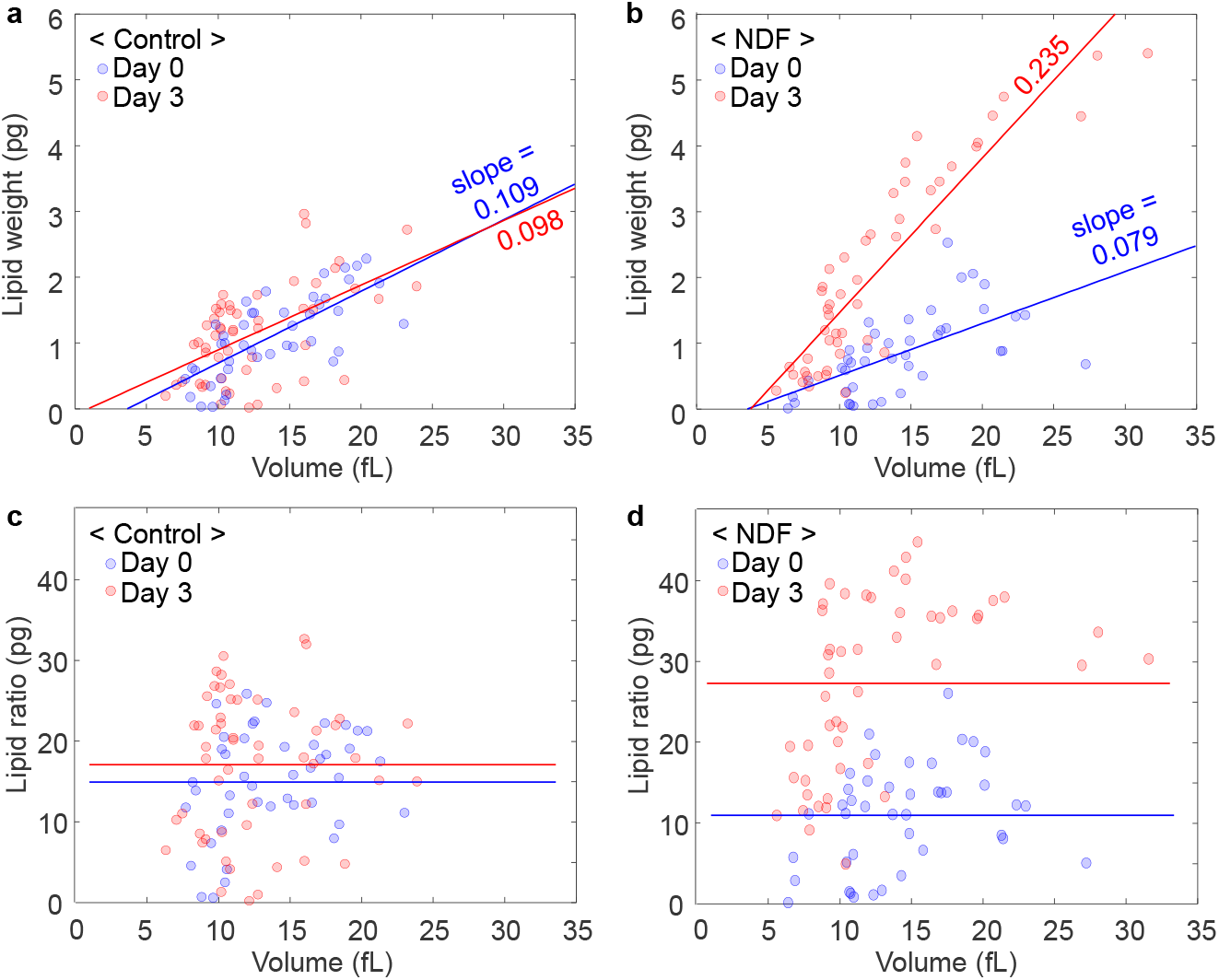
Scatter plots of the lipid weights (a-b) and lipid ratios (c-d) according to the volume of individual *N. oculata* cells. The circles represent the individual values measured by ODT. (a) The correlation coefficients between the lipid weights and volumes are 0.73 and 0.56 for Day 0 and Day 3, respectively. (b) The correlation coefficients between the lipid weights and volumes are 0.62 and 0.90 for Day 0 and Day 3, respectively. (c) In the control group, the correlation coefficients between the lipid ratios and volume are 0.31 and 0.07 for Day 0 and Day 3, respectively. (d) In the NDF group, the correlation coefficients between the lipid ratio and volume are 0.32 and 0.48 for Day 0 and Day 3, respectively. The solid lines represent results of linear fitting in (a-b) and the average lipid ratios in (c-d), respectively.

In addition, the relationship between the lipid ratio and cell size was investigated to answer how the lipid ratio changes with the cell volume (Fig. 6c and 6d). The results show broad distributions of the lipid ratio around the average values (indicated as solid lines in Fig. 6c and 6d). Again, we observe that the average lipid ratio increases in the NDF group after cultivation while the control group shows the considerable overlap of the data from Day 0 to Day 3. The important result is that the correlation coefficients between the volume and lipid ratio were found to be 0.19 and 0.40 (on average) for the control and NDF groups, respectively. These weakly positive coefficients appear to suggest that there is no strong relationship between the lipid ratio and cell size.

## Discussion

We presented the label-free, non-invasive, quantitative method to measure lipid contents in microalgal cells using 3D RI tomography. We measured the 3D RI distribution of the individual living cells to identify and quantify lipid accumulation. The identification of the lipids using the RI threshold was verified with fluorescence imaging using Nile red dye. The measurements of size, DCW, and lipid content of *N. oculata* cells showed agreement with conventional approaches. Effects of nitrogen deprivation on cell lipids were quantitatively investigated, and related parameters were investigated using correlative analysis. We believe ODT has potential for lipid quantification and biological studies of living cells.

In addition to the capability for label-free quantitative 3D imaging, another advantage of ODT is that it does not require illumination with strong light. This is particularly important for studying organisms such as microalgae because the strong light disturbs the metabolism of photosynthetic organisms. In this work, the light power measured at the sample plane was 8 μW, while 20 μW is sufficient for recording holograms at 100 Hz. Because the intensity was distributed over the whole field of view (approximately 100×100 μm^2^) rather than focused on a spot, the effects of the illumination on the sample are further reduced compared with other techniques using point excitation. The unique advantages of ODT: label-free imaging, quantitative RI contrast, and minimal light exposure, enable imaging and investigating live microalgae in a culture medium without disturbing their cell physiology.

From the technical point of view, additional optical properties could be utilized to improve the molecular specificity. For example, we expect that polarization-sensitive holographic imaging^51^ could be utilized to identify starch-granule reserved in the microalgae because the starch granules exhibit strong birefringence. Spectroscopic modality could also be employed in ODT to exploit RI dispersion^52, 53, 54^. Because the dispersion also reflects the unique wavelength-dependent optical properties of materials, more components in the microalgae could be quantified if higher specificity and greater sensitivity were provided.

For more practical application of ODT in lipid-quantification, high-throughput imaging would be beneficial. The current throughput of ODT imaging is limited to the measurements of a few cells per minute including acquisition time and manual translation of a sample stage to find cells. High-throughput imaging could be achieved by reducing the acquisition time and by utilizing a motorized translational stage. The total acquisition time for one 3D distribution (two seconds in this work) could be further reduced (to less than 0.1 s) by optimizing the number of 2D holograms^55^. Simultaneous 3D fluorescence imaging with several fluorescence channels would also enhance the capability for correlative analysis^56^, and this is already commercially available. The mechanical translation of a sample stage could also be replaced by a fluid flow system. In the future, we believe that real-time *in-situ* monitoring of lipid productivity could be provided by dynamic ODT via fluidic channels linked to culture systems.

## Materials and Methods

### Algae cultivation

A sample of the micro-algal species *N. oculata* (UTEX LB2164) was obtained from the Culture Collection of Algae at the University of Texas at Austin. The cells were grown under continuous illumination by fluorescent lamps at 100 μmol·m^-2^·s^-1^ and aerated continuously with 2% CO_2_-balanced air at a flow rate of 40 mL/min. The cultures were grown in 0.5 L bubble-column photobioreactors containing 0.4 L of modified f/2 medium at 20 °C. The cells were harvested in the middle of the exponential growth phase (2.0 ± 0.3 g fresh weight/L), and collected by centrifugation. The collected cells were washed twice using fresh medium without nitrogen to remove nitrogen sources. The washed cells were re-suspended in the fresh medium without nitrogen and split equally into two cultures. One culture was the NDF group, and the other was supplied with nitrogen (NaNO_3_, 225 mg/L) as another control group. These cultures were grown for three days.

### Optical diffraction tomography

The 3D RI distribution of individual algae was reconstructed from multiple 2D optical fields measured at various illumination angles. The light diffracted from a cell was recorded as a spatially modulated hologram using a quantitative phase imaging technique^18, 57^. The 2D optical field at the sample plane was quantitatively retrieved from the hologram via a field retrieval algorithm^58, 59^. From the retrieved optical fields, the 3D RI distribution within the cell was reconstructed according to Fourier diffraction theorem^22, 60^. Holograms (201 of them) were measured at various illumination angles to reconstruct each 3D RI tomogram. The maximum incident angle was equivalent to 45° at the sample medium. The missing information from limited illumination and detection angles of the optical lenses was filled using an iterative non-negativity constraint^61^. Visualization of the 3D iso-surface was conducted using commercial software (TomoStudio™, Tomocube Inc., Republic of Korea).

In order to measure the 3D RI tomograms of individual algae, we utilized a commercial ODT system (HT-1H, Tomocube Inc., Republic of Korea) with modifications for fluorescence imaging. The ODT system used, is based on Mach-Zehnder interferometry, as shown in Fig. 1. A diode-pumped solid-state laser (532 nm in vacuum; Samba™, Cobolt Inc., Sweden) was used as the light source. The laser beam was split into two arms, a sample beam and a reference beam, using a 1×2 fiber coupler. A digital micro mirror device (DMD), placed on the plane conjugated to the sample, was used to control the illumination angle ^62^. With this system, a plane wave with controlled illumination angle impinges on a sample via a high numerical aperture (NA) condenser lens (NA = 1.2, water immersion, UPLSAPO 60XW, Olympus Inc.) and is detected via a high NA objective lens (NA = 1.42, oil immersion, PLAPON 60XO, Olympus Inc.). The sample beam and reference beam are combined using a beam splitter to generate a spatially modulated hologram at the camera plane, which is recorded by a CMOS camera (FL3-U3-13Y3M-C, Point Grey Research Inc.). The total acquisition time for 201 holograms was approximately 2 s (10 ms per hologram). The theoretical spatial resolution of the system was 119 nm and 336 nm for lateral and axial directions, respectively^36^.

### Wide-field fluorescence imaging

To validate identification of the lipids in *N. oculata*, wide-field fluorescence imaging with Nile red dye was used. The same optical imaging system was used for both ODT and fluorescence measurements. The same laser was used as an excitation light source for the Nile red because the excitation spectrum of the dye includes the wavelength of the laser. Once the ODT measurements of a sample were finished, a notch filter with a central stopband wavelength of 532 nm (FL532-1, Thorlabs Inc., USA) was manually inserted in front of the camera to block the excitation laser light. Then, wide-field fluorescence images of the cell were captured with an exposure time of 800 ms.

### Algae cell preparation

For single-cell imaging, *N. oculata* solution taken from a culture bath was diluted with fresh medium so that the cell density became approximately 10^6^ cells/mL. For fluorescence staining of the lipid inside cells, Nile red (19123, Sigma-Aldrich Inc.) solution in dimethyl sulfoxide (10 mg/mL) was added to the diluted sample solution such that the final concentration of Nile red in the solution was 5 μg/mL. The sample solution was then incubated at room temperature in darkness for 3 min. Then 5 μL of the sample solution was sandwiched between a pair of coverslips. Double-sided tape 100 μm thick was inserted between the two coverslips as a spacer.

### Emulation of fluorescence-like images

To generate fluorescence-like images from the measured 3D RI tomograms, we applied 3D convolution on the identified high RI regions (*n* > 1.46) using a 3D point spread function (PSF) of the wide-field microscopy. The 3D PSF was numerically generated based on NA = 1.42 and 630 nm wavelength. This imaging processing emulated the situation in which the lipid droplets were uniformly stained with Nile red dye.

### Quantification of the cell concentration and size

Cell concentration and cell size distribution were measured using a Coulter Counter (Multisizer 4, Beckman Coulter Inc., Brea, CA, USA). Fresh weight was determined by the data from the Coulter Counter for microalgal growth^63^. Cell size distribution data were used for comparison with the cell volumes of 3D RI distributions.

### Quantification of the dry cell weight

To measure the dry cell weight (DCW), 10 mL of each culture was collected, centrifuged; then dried in an oven at 80 °C for 12 h. The dried weight was measured using an analytical balance (HR-250AZ pharma balance, A&D Weighing, USA) and DCW (g/L) was calculated from the dried weight and initial volume of the culture sample.

### Quantification of the fatty acid content

Fatty acid transesterification was conducted to analyze the fatty acid content of freeze-dried *N. oculata* cells. The cells were collected using centrifugation, and then washed twice with distilled water. Fatty acids were extracted in a mixture of acetyl chloride and methanol (5:100, v/v) with methyl nonadecanoate as the internal standard, at 80 °C for 1 h in a drying oven. The fatty acid analysis was conducted using a gas chromatograph (YL6500GC, Young Lin Inc., Anyang, Korea) with a flame ionization detector, capillary column (HP-INNOWAX, 30 m and 0.53 mm internal diameter), and helium as the carrier gas.

## Acknowledgement

This research was supported by KAIST, BK21+ program, Tomocube, and National Research Foundation of Korea (2015R1A3A2066550, 2014M3C1A3052567, 2014K1A3A1A09063027, 2016M1A5A1027462), and the Marine Biotechnology Program (20090267) funded by the Ministry of Oceans and Fisheries, Korea.

## Author Contributions

D.-J.K, C.-G.L. and Y.P. conceived the idea and directed the work. J.J., G.K., and M.L. performed experiments and analyzed data. S.-J.H. and H.-B.K. prepared the algae samples and analyzed the sample using gas chromatography. S.S. and S.L. provided analysis tools and analyzed data. All the authors wrote the manuscript.

